# Towards a real-world brain-computer interface for image retrieval

**DOI:** 10.1101/576983

**Authors:** Ben McCartney, Jesus Martinez-del-Rincon, Barry Devereux, Brian Murphy

## Abstract

Brain decoding — the process of inferring a person’s momentary cognitive state from their brain activity — has enormous potential in the field of human-computer interaction. In this study we propose a zero-shot EEG-to-image brain decoding approach which makes use of state-of-the-art EEG preprocessing and feature selection methods, and which maps EEG activity to biologically inspired computer vision and linguistic models. We apply this approach to solve the problem of identifying viewed images from recorded brain activity in a reliable and scalable way. We demonstrate competitive decoding accuracies across two EEG datasets, using a zero-shot learning framework more applicable to real-world image retrieval than traditional classification techniques.

## Introduction

Research in the field of Brain-Computer Interfaces (BCI) began in the 1970s [1] with the aim of providing a new, intuitive, and rich method of communication between computer systems and their users. Typically, these methods involve measuring some aspect of neural activity and inferring or decoding an intended action or particular characteristic of the user’s cognitive state. Although BCI is still in its infancy, there are already practical applications in assistive technology as well as disease diagnosis [2, 3]. Brain-controlled prosthetics [4] and spellers [5] have shown their potential to enable natural interaction in comparison with more traditional methods, such as mechanical prosthetics or eye-movement-based spellers. Other relevant applications include identifying the image that a user is viewing, usually referred as image retrieval, of particular interest in the field of visual attention applied to advertising and marketing, searching and organising large collections of images, or reducing distractions during driving, to name a few.

Although brain decoding technology has immense potential in diverse applications, it faces multiple challenges related to speed and accuracy that must be overcome before it emerges as a disruptive technology. The complexity of BCI stems from the naturally low signal-to-noise ratio (SNR) and high dimensionality of raw brain data, which often complicates automated analysis and can force researchers to manually analyse previously recorded neural activation data. This is typically done either by examining the frequency domain or by plotting Event-Related Potentials (ERPs). In an ERP experiment the participant will be presented several times with a similar stimulus and their neural response each time can be recorded and averaged. These ERPs can be analysed against well-known response patterns or, alternatively, characteristics such as the strength and timing of signal peaks can be quantified and analysed automatically. ERP analysis is well established and has strong applications in medical diagnosis [6] and in cognitive neuroscience research [7, 8]; however, the broad characterisation of brain response used in traditional ERP methods is not richly informative enough to decode the level of detail required to make predictions about a participant’s cognitive state, as required for BCI image identification.

Given the complexities of decoding the nature of an arbitrary visual stimulus from a person’s brain activity, cognitive neuroscientists and BCI researchers have traditionally tackled the simpler task of determining which of some finite set of category labels corresponds to a particular pattern of brain activity. In one of the first such studies, Haxby and colleagues [9] collected functional Magnetic Resonance Imaging (fMRI) data as participants viewed a series of images from the categories of *human faces, cats, houses, chairs, scissors, shoes* and *bottles*, along with images of *random noise*. The researchers were able to determine with 83% accuracy which category of object the participant was viewing.

However, fMRI is impractical for general BCI applications. Murphy et al. [10] used Electroencephalography (EEG) rather than fMRI and achieved 72% accuracy in classification across the two classes of mammals and tools. While this study addressed a much simpler problem with only two possible classes, it demonstrated category decoding using relatively inexpensive and less intrusive EEG data collection methods (fMRI and EEG technologies are discussed in more detail in Section ‘Brain Data’).

In the studies mentioned above the classifiers would not determine specifically which stimulus image was displayed (as required for image retrieval), instead they only determine the category which the stimulus image belongs to. Moreover, as a classification approach, this is not scalable to new classes and, although it may yield a high accuracy, it becomes less accurate with increasing number of classes. An alternative approach to BCI image classification makes use of rapid serial visual presentation (RSVP) [11, 12]. The participant is presented with a rapid stream of images (approximately 10 each second) and is instructed to count the number of times a particular target image or object appears. A classifier can then reliably decode whether for a given segment of brain data, the participant had been presented with a target or non-target image. This RSVP approach could be more directly applied to our problem by showing a participant a target image from a gallery, and then presenting all of the images in a gallery one by one with the expectation that when the target image should illicit neural activity sufficiently different from the non-target images to identify it. However, as the number of images in the gallery grows, it becomes impractical to present them in a real-world searching scenario.

As a more scalable solution, zero-shot learning presents a novel approach to brain decoding in which some feature space is created which can describe each stimulus class, and a mapping is defined between neural activation data and the stimulus feature space. Such a mapping can be defined with a subset of the full set of classes and/or instances, and tested using withheld classes/instances. With this approach, the system can decode arbitrary stimulus images it has not yet been exposed to. Introducing a feature-based model comes at a cost however, as it also impacts the overall accuracy of the system. At present most zero-shot systems in this area [13, 14] exhibit performance insufficient for real-world applications.

Following the work in Haxby et al. [9], Mitchell et al. [13] used fMRI to decode the meanings of nouns corresponding to concrete objects, using as features each noun’s textual co-occurrence frequency with a set of 25 verbs. This study was one of the first to make use of zero-shot learning, allowing them to decode classes (i.e. nouns) from outside the training set. Others have used visual image features rather than semantic features to decode cognitive states associated with viewing stimulus images in a zero-shot framework [15]. Using a Gabor-based voxel decoder, Kay et al. [15] achieved accuracy of 51% and 32% among 1620 images in a single-trial identification task, for 2 distinct subjects. These and other studies, while using fMRI rather than EEG data, demonstrate the relevance of both semantic and visual information in image decoding.

In related work, Palatucci et al. [14] used a similar procedure to Mitchell et al. [13] but with EEG data. In a study by Carlson et al. [16] using visual images, Linear Discriminant Analysis (LDA) was used to determine the categories of objects presented to participants. The aim of this study was to map out the stages of object recognition by comparing the decoding accuracies across different levels of object representation and different time windows. They found that the peak decoding rate for distinguishing between images of the human body was 120ms after the image appeared, whereas the higher-level semantic distinction between animate and inanimate images was best determined after 240ms. Using a combination of low-level visual features and semantic features, Clarke et al. [17] demonstrated that decoding accuracy was significantly improved by the incorporation of the semantic features from around 200ms post-stimulus-onset. Similarly, Sudre et al. [18] also obtained high decoding accuracy using brain data using both visual and semantic feature sets. These studies both use magnetoencephalography (MEG), which is impractical for real-world BCI technology; however, their conclusions suggest that decoding of neural activation data using visual and semantic models can be a feasible approach to image decoding in a real-world BCI framework.

In this project, we aim to make use of these different levels of information in brain activity explicitly by using specifically chosen feature generation models rather than implicitly by grouping our images into different categories. We also aim to perform a more difficult task: where Carlson et al. [16] utilises zero-shot learning only to determine membership of the stimulus to a particular object category, our approach will aim to determine which actual image was viewed. To this aim, this paper proposes an EEG zero-shot learning framework for individual image retrieval which makes full use of both advanced visual and semantic image features. This approach is motivated by the ultimate goal of designing a system which can retrieve any arbitrary image specified by a neural activation generated by a user thinking about that image, although as a preliminary step towards this goal we restrict our experiments to cases where images are viewed rather than imagined.

The main contributions of this paper are:

- First time visual and semantic features are used together for EEG zero-shot learning, which translates to potential for a real-world BCI image retrieval system.
- State-of-the-art performance for the particular task of EEG-driven image retrieval in a zero-shot framework.
- Evaluation across two datasets from different sources including a large open dataset for future comparative studies.
- Analysis of how well the feature sets chosen reflect the expected brain activity.

## General Methodology

Our framework comprises of three main components. First the brain data must be cleaned and a subset of the EEG features extracted to represent the underlying cognitive states. Then we apply our chosen computer vision and semantic models to the stimuli, to create a representation of each image in this visuo-semantic feature space. Finally we use a linear regression algorithm to find a mapping between the brain and stimulus spaces which makes the brain decoding possible. A high-level overview of this architecture can be found in Figure 1.

**Fig 1.**
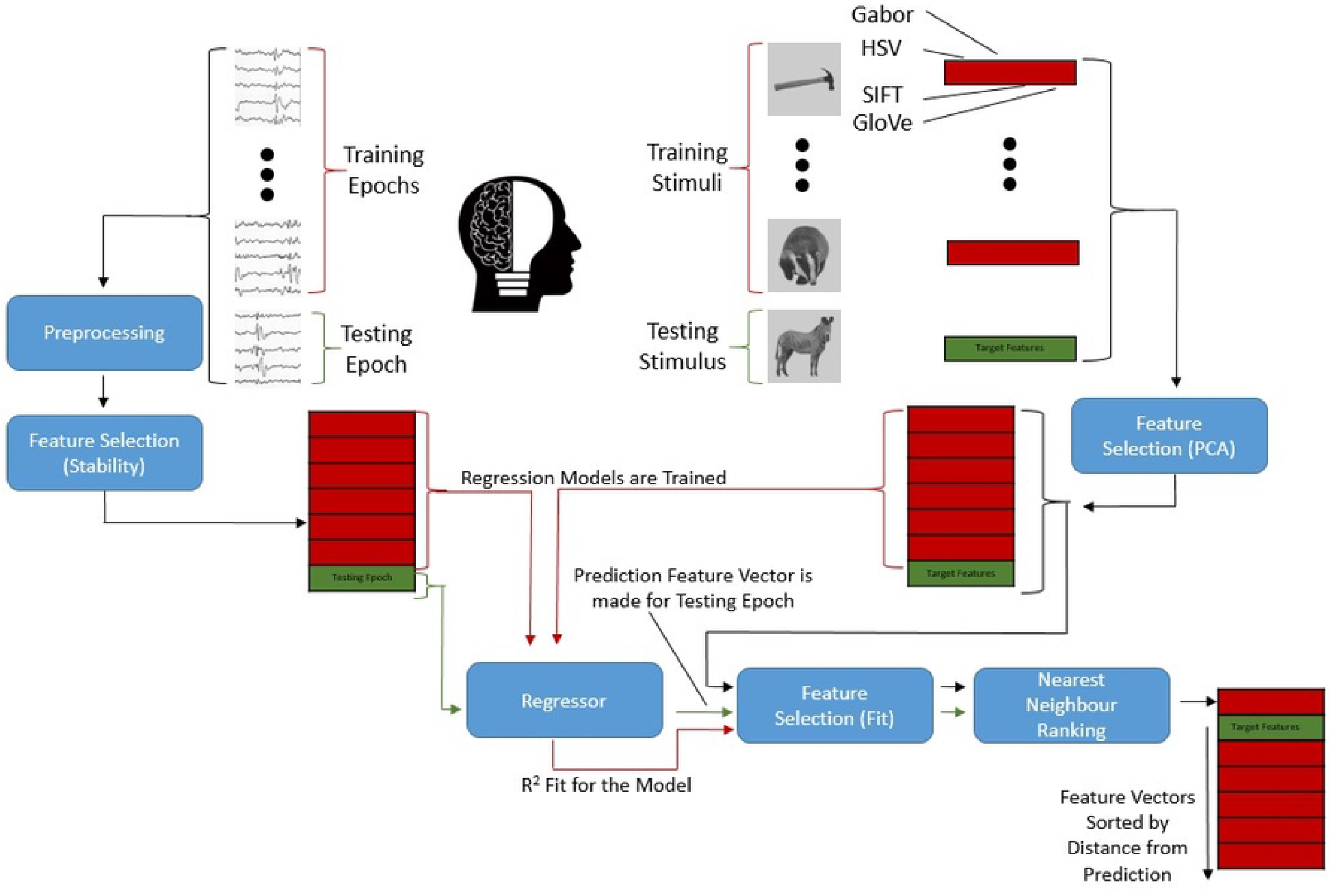
Information flow in the image retrieval architecture. Overview of the flow of information and processing during a single fold of cross-validation (‘Zero-shot Prediction’ Section). Model performance is determined by fit of the predicted feature vectors: in the example above, the true target features are in the second position of a sorted list of neighbours. In this case with a total of seven possible images, this results in a rank of 2, and a CMC AUC of 78.57% (‘Measure of Accuracy’ Section).

### Brain Data

Two of the most widely used approaches to recording brain activity are functional Magnetic Resonance Imaging (fMRI) and Electroencephalography (EEG). The former can localize the physical source of brain activity with high spatial accuracy. However, the temporal resolution of fMRI is limited to a sampling rate of 1-2 seconds. Moreover, fMRI requires an MRI scanner, a large and expensive piece of equipment using powerful magnetic fields and liquid helium coolant, making it unsuitable for BCI systems outside of laboratory or clinical settings. As a cheaper and more convenient alternative, Electroencephalography (EEG) can be used to measure the electrical activity produced in the brain. As neurons communicate, they produce a small electrical current. Individually this electrical activity is weak, however, often these cells fire in groups and produce an effect strong enough to be detected by sensitive conductors (or sensors) placed around the scalp. EEG is the technique of recording these small variations in electrical activity as a multi-channel signal. Measuring the electrical activity rather than blood oxygenation (as in fMRI) gives much higher temporal resolution, and as such EEG can operate in windows of <10ms. However, the collective nature of the electrical activity detected by EEG – and the fact that the electrical activity must propagate to the scalp – make it much more challenging to localise each source. There is therefore a trade-off between cost/convenience and the quality of the information recorded; compared with fMRI (or MEG), EEG data is easier to obtain, but is more difficult to analyse in terms of the underlying brain activity. EEG is also impacted by a greater sensitivity to a variety of external artefacts, such as muscle movement, cardiac activity, ambient electrical activity, and electro-ocular activity, all of which negatively impact SNR. Some of these noise sources can be isolated and removed either with signal processing algorithms or by hand.

As we are interested in eliciting cognitive states associated with particular images, the experimental paradigms used for the EEG data in this study involve repeated presentations of images on a computer screen (‘Datasets’ Section). Each time an image is presented is termed a “trial” and the small window of EEG data associated with these trials are known as “epochs”. We use epochs which begin when an image is presented and are one second long to comfortably encompass the informative brain activity [16]. It is these epochs which we attempt to map to images features and aim to determine which image was presented at the time the epoch was recorded.

### Preprocessing Approach (FASTER)

Preprocessing is a necessary stage of EEG data analysis that involves aligning, normalising and otherwise cleaning the raw data in order to make it more suitable for downstream analyses. The main goal of preprocessing the EEG data in our framework is to remove sources of noise in order to minimise obfuscation of underlying useful patterns in the data. Recordings are first filtered to remove ambient interference. One of the strongest noise sources in EEG is ambient electrical activity near the recording equipment, such as personal computers, large lights, or improperly insulated wiring. These signals are relatively easy to separate from brain activity based on their frequency, typically 60hz or 50hz (in America and Europe respectively). A lower frequency cut-off must also be established to remove slower sources of noise – these are generally slow changes in the electrical profile of the scalp or sensors such as a gradual increase or decrease of perspiration leading to a change in conductivity. A band-pass filter was used to remove any signals in our data with a frequency outside the range 1-40hz as in [10, 11, 14, 19–22].

**Fig 2.**
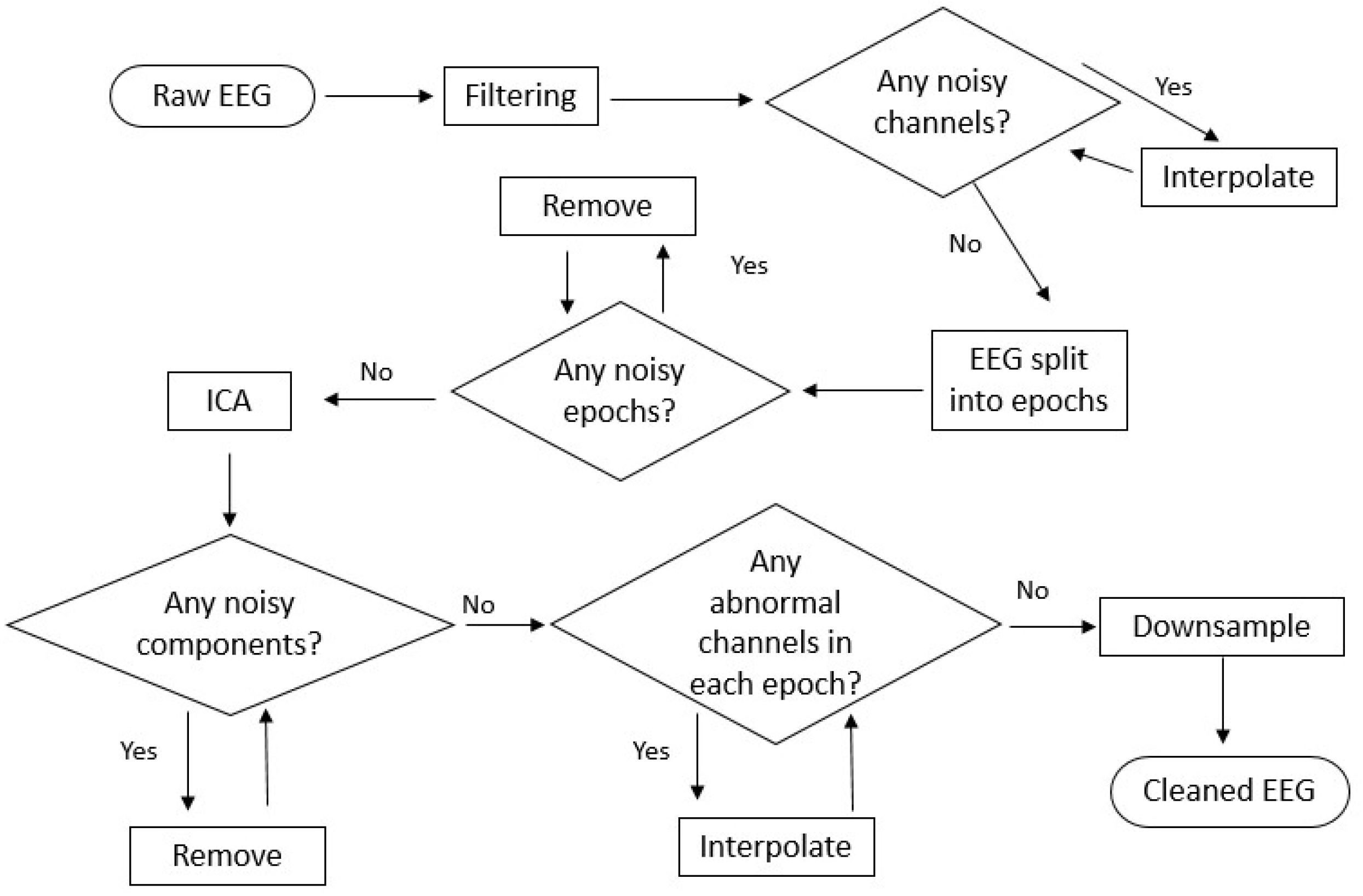
Preprocessing Overview.

Channels with poor contact with the scalp were then identified using the variation, mean correlation and Hurst component, and these were removed and then interpolated from nearby sensors similar to [19, 23]. A section of the EEG lasting one second was extracted each time an image was displayed. These epochs were baselined [19, 22] by subtracting the average value from the 500ms prior to the image presentation. Epochs which fell outside the threshold for amplitude range, variance or channel deviation were removed as in [11, 19, 21, 23]. Following this, Independent Component Analysis (ICA) [24] was performed primarily to identify artefacts related to eye movement as in [10, 19, 20]. In this step the input signal is decomposed into an approximation of its sources, each component is then correlated with sensors placed nearest the eyes and thresholds set for spatial kurtosis, Hurst exponent and mean gradient. Components identified as isolating sources of noise are removed and the EEG signal reconstructed from the remaining components.

Next, within each epoch, channels were examined for short term artefacts using variance, median gradient, amplitude range and channel deviation. Channels identified as noisy within the bounds of the epoch were replaced by an interpolation from other nearby channels within that epoch. The recording is also downsampled to a rate of 120hz as in [11, 14, 20] to reduce dimensionality before machine learning is applied.

All the above preprocessing steps were implemented using the EEG preprocessing toolkit FASTER [19].

As a final preprocessing step before the EEG data are used in our regression model, the data are z-scored (standardised). We primarily perform this step to ensure that the mean of the data is zero as this can simplify the parametrisation of our machine learning. This takes place each iteration of the cross validation, the mean values for the transformation are calculated using only the training samples and the transformation is then applied to the training and testing samples to avoid any influence of the latter in the former.

### EEG Feature Selection

After preprocessing, an EEG feature extraction process is used to continue reducing the dimensionality of the data by extracting the most discriminatory features from the preprocessed data, and further removing uninformative and noisy dimensions of the data. This facilitates the successful mapping of EEG data to our image feature space by extracting only those aspects of the EEG signal which are likely to be informative about the visual and semantic feature sets. Following the approaches used in Mitchell et al. [13] and evaluated in Caceres et al. [25], we ignore all but the features with the highest collinearity across presentations of the same stimulus on the screen. Concretely, the EEG data for a particular participant following preprocessing is a 3D-matrix of size nE × nC × nT, where nE is the number of epochs (i.e. the number of stimulus presentation events), nC is the number of channels (or sensors) in the EEG headset, and nT is the number of timepoints in an epoch (the number of times during an epoch sensor values were recorded). In this work, we use an epoch length of one second and downsample the data to 120Hz, giving nT = 120. We treat the data from each time sample and each sensor as a separate feature, giving a total of nC × nT candidate features. In order to calculate feature collinearity, we reshape the nE × nC × nT data matrix to a 2D-matrix of size nE × (nC × nT), or, equivalently, (nS × nP) × nF where nS is the number of stimuli, nP is the number of times each stimulus was presented in a recording, and nF is the number of EEG features. We then transform this back into a 3D-matrix of shape nF × nP × nS and term this matrix *D. D* is therefore composed of a nP × nS feature matrix for each EEG feature *f*. To calculate a stability score for a feature, we measure the consistency of the feature across different presentations of the same stimulus – we calculate the Pearson correlation for each pair of rows in D and use the mean of these correlations as the stability score for that EEG feature *f*:

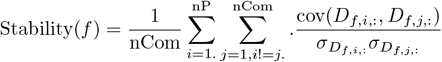

where *σ_x_* is the standard deviation of *x* and

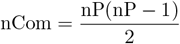

In each iteration of our cross-validation, we calculate the stability of each EEG feature using the training set and select the most stable features for fitting the regression model. More detail of the cross-validation architecture can be found in the ‘Zero-shot Prediction’ Section.

### Image Feature Space

Computer vision is the field of study devoted to designing algorithms to interpret digital images, so it is a natural place to look for a feature space. Different computer vision models extract features at different levels of abstraction, ranging from recognising simple lines or colours through to recognising objects. Previous research [16, 17, 26–29] shows that these levels of abstraction are evaluated sequentially in the human ventral visual processing stream. In light of these findings, we expect earlier EEG features to contain predominantly low-level visual information, with higher-level visual features being increasingly present in later EEG features. For maximal decoding performance, it is therefore essential to find a set of computer vision models which cover each level of abstraction that will be represented in the EEG features. Furthermore, we chose feature sets which are grounded in similar mechanics to human visual processing, under the rationale that these feature sets have the potential to best match with human brain activity.

### Gabor Filters

In order to model human edge and texture detection we chose to use Gabor Filter Banks as this well-established computer vision technique identifies visual edges in a very similar way to the lowest-level of human visual processing in cortical areas V1 and V2 [30–33]. A bank is comprised of a set of filters which each represent an edge with a particular orientation and spatial frequency, these filters can be used to identify where in an image there is a matching edge. The filter bank used here contains eight evenly spaced orientations (*θ*) and four standard deviation values (*σ*) ranging from two to five, resulting in a bank of 32 filters. The rest of the parameters were fixed at default with ksize = (31, 31), wavelength of the sinusoidal factor (*λ*) = 6.0, spatial aspect ratio (*γ*) = 0.5 and phase offset (*ψ*) = 0.

Each pixel co-ordinate in an image *x, y* is convolved with a Gabor filter described by the parameters above:

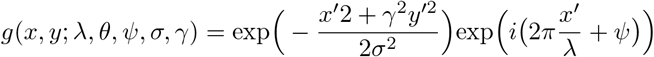

where

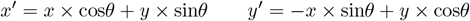

Let Lϑ and Lσ denote the sets of parameter values defining the filter bank:

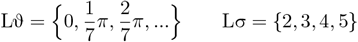

Each image in our feature set was convolved with every filter, and the result summed to generate a histogram of 32 dimensions *v_gabor_* for each image:

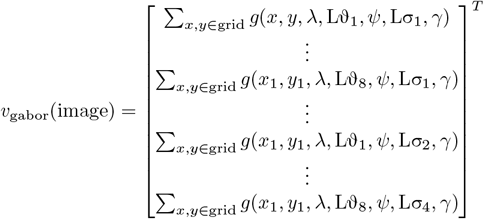

We stack the *v*_gabor_ vectors to create the final matrix of Gabor features for our image set:

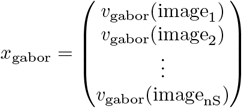

### Scale Invariant Feature Transform

The brain is also sensitive to higher-level visual information which is not adequately captured by simple and spatially local Gabor Filters. In order to make use of higher-level visual processing in our system we chose to apply a prominent computer vision model which detects and describes keypoints in an image; locations with particular visual saliency. Scale Invariant Feature Transform (SIFT) identifies these keypoints using difference of Gaussians and generates a descriptor for the pixel neighbourhood [34]. We then applied Visual Bag Of Words (VBOW) [35] to the extracted SIFT features. Implementing VBOW involves generating SIFT keypoint descriptors for a large corpus of images and then selecting the most informative descriptors to compile a codebook. Our goal in this project is to create an approach applicable to real-world BCI systems, and to achieve this our image feature space must have some capacity to describe and discriminate novel images. To this end, we used a SIFT codebook trained with a large, diverse corpus of images taken from ImageNet [36].

Each SIFT Descriptor represents a 4 × 4 grid around some keypoint in the image. Difference of Gaussian (DoG) is run on each segment of this grid and the results compiled into a histogram with eight bins. A SIFT descriptor is the resulting 8 × 4 × 4 = 128 dimensional vector, indexed by *x-y*. Each element of the codebook is a SIFT Descriptor. K-means clustering was performed with a random subset of 10 million SIFT descriptors generated over the ImageNet corpus to produce 1000 clusters. The centroid of each cluster was then taken to produce a codebook of 1000 dimensions to categorise future SIFT descriptors.

Using VBOW has the benefit of finding features that generalise well across multiple different objects and as such have the best chance of extending to new classes. Moreover, it removes spatial data making the feature vector invariant to spatial transformations such as rotation, translation and scale which is less relevant to intermediate-level visual information. A list of imageDescriptors were generated for an image, and used to produce a histogram *v*_shift_ of how often each ‘visual word’ encoded in the codebook appeared in the stimulus image.

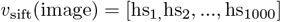

where

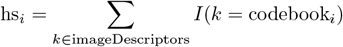

and *I* is an indicator function evaluating to 1 if the argument is true and 0 otherwise.

This implementation made use of Dense SIFT, meaning the keypoints correspond to a regularly sampled grid, rather than a set of natural keypoints estimated for an image. A histogram *v*_sift_ was generated for each image, and collated into a matrix representing our stimulus image SIFT features *x*_sift_.

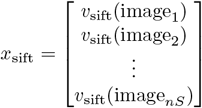

### Colour Histogram

Finally, as none of the previous visual features encapsulates colour information, we chose a global HSV histogram to model colour in our approach, since there is some evidence that a HSV colour space comes closer to reflecting human vision than RGB [37]. A HSV histogram *v*_hsv_ is generated for each image using a quantisation of four bits per pixel and channel:

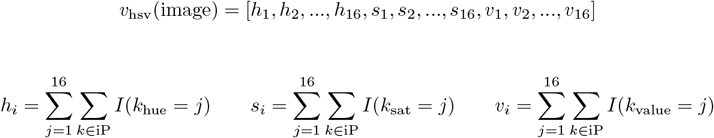

Where iP is the list of pixels in the image, and *k*_hue_, *k*_sat_ and *k*_value_ are the hue, saturation, and value of the pixel *k* respectively. This gives each HSV channel 16 bins to produce a histogram of 48 features. The histograms are then collated into a matrix representing our HSV feature space *x*_hsv_.

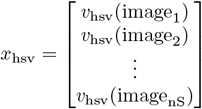

### Global vectors for Word Representation

While visual features allow us to describe images at low and intermediate levels, higher-level semantic processing requires us to characterise the image in terms of the object it contains. To model object-level information, we included a general set of features describing the semantic differences between concepts. We chose a set derived with the Global Vectors for Word Representation (GloVe) algorithm as it is well established [38–40] and state-of-the-art vector datasets are readily available. The learning objective of GloVe is to generate for each word an *M*-dimensional vector such that the dot product of two of the vectors equals the logarithm of the probability of the associated words co-occurring in text. As *M* increases, a larger number of words can be more accurately described; however this increases the computation time both in training GloVe and in any downstream analyses which use GloVe vectors as input. In this project, we make use of a pretrained matrix gMat of 1.9 million words with 300 dimensions indexed by the word.

Firstly a name is assigned to each stimulus image to describe the subject of the image. A number of our stimulus images were labeled with a Multi-Word Expression (MWE) which did not have a corresponding feature vector in *gMat*. In these cases we used the mean of its composite words, following [41]. For example, the stimulus “plaster trowel” was set to the mean of the vector for “plaster” and the vector for “trowel”.

For each of our images we chose a single word or MWE to represent the content (i.e. the depicted object), and take the row of the GloVe matrix which corresponds to that word as the feature vector for the image in our high-level semantic feature space.

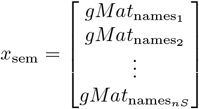

where

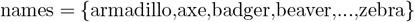

### Combining the Feature Sets

The complete visuo-semantic feature set is then composed by combining *x*_gabor_, *x*_sift_, *x*_hsv_ and *x*_sem_. Concatenating the raw feature sets together would result in a poor and imbalanced feature space due to the differences in dimensionality and value scaling across the different constituent feature sets. We therefore normalise each feature set to ensure that the values in each row range from zero to one and perform Principal Component Analysis (PCA) to reduce the dimensionality of the concatenated feature space.

With *y* ∈ *x*_gabor_ and *z* be the length of *y*, we normalise the feature vector by its range and stack the results to form *x*_gaborR_:

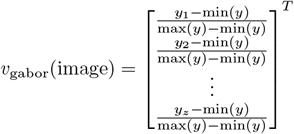

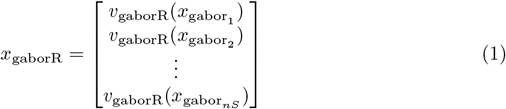

The final complete set of features is the concatenation of the features from each of the component visual and semantic models:

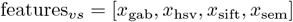

Finally, before using these features in our classification model, we apply one further feature selection based on a measure of fit from the regression model (as described in Section ‘Zero-shot Prediction’).

### EEG Mapping

The mapping of our EEG data to our visuo-semantic feature space is essentially a problem of fitting a regression model *f_i_* for each image feature *i* such that *f*(*EEG_y_*) = [*f*_1_, …,*f_i_*] ≈ features_*y*_ where EEG_*y*_ is the preprocessed nC × nT-dimensional brain activity vector associated with stimulus image *y*. Assuming a linear relationship exists between these two components, multiple linear regression can be applied to find some set of weights w1 such that *f*_1_(EEG_*y*_) = *v*_EEG_1__ * w1_0_ + *v*_EEG_2__ * w1_1_ + … will produce a value as close as possible to features_*y*_1__, some vector of weights w2 such that *f*_2_(EEG_*y*_) = *v*_EEG_1__ * w2_0_ + *v*_EEG_2__ * w2_1_ + … will produce a value as close as possible to features_*y*_2__, and so on until a vector can be stacked which is as close as possible to features_*y*_.

Prior studies [10, 13, 14] have shown success using a linear regression model with brain data when they are regularised. This coupled with its speed and simplicity made it a natural choice for a baseline approach. L2 regularisation is used to reduce overfitting and improve the generalisation properties of the model. This choice is preferred over L1 regularisation given the expected high collinearity of our samples, i.e. signals recorded from nearby locations in very similar temporal instants should register very similar sources in brain activity. A good model will be able to generalise the relationship rather than being limited to projecting the particular samples and/or classes used in training. If this is achieved, the mapping mechanism and the representative feature spaces could be used within a zero-shot learning architecture.

### Zero-shot Prediction

Once a mapping between EEG data and the image feature space has been learned from training, a prediction of image features can be made for an EEG epoch withheld from the training set. To ensure a zero-shot framework, we use leave-one-class-out cross-validation to iteratively withhold all epochs associated with a particular stimulus/image for testing in each iteration. Concretely, this means we withhold the data for trials related to Stimulus 1, and train a regression model from the trials for the rest of the stimuli. We then pass the withheld testing trials into our regression model to produce a predicted image feature vector for each trial. We then return the trials for Stimulus 1 to the training set and instead withhold the trials for Stimulus 2. A separate regression model is trained from scratch for this new training set, and then predicted image feature vectors are produced for the Stimulus 2 trials. This pattern is repeated for each stimulus image in the recording.

Following the regression, there is one final step of feature selection over the predicted image features before moving to the feature matching for image retrieval. We do not make use of all image features in the predicted image feature vector, but instead select just those which are best represented in the EEG data. To make the distinction between useful and under-represented features, we approximate each feature’s informativeness by calculating the measure of fit of our regression model. When predictions are fed to the classifier, we ignore the columns of the feature space and the predicted feature vectors with the lowest measure of fit. For each iteration of train/test split, after the regression model has been fit, an R^2^ measure of fit is calculated for each image feature column in features. For each epoch in a recording we produce a predicted image feature vector and collate these vectors into the matrix *p*. Each epoch is associated with a particular stimulus image and each stimulus image is associated with a feature vector in features, so we generate *t* such that *t_i_* is the feature vector associated with the stimulus image used in epoch *i*.

These fit values are then averaged across iterations to produce an estimate of which image features are best represented in the EEG data. This estimation is reached entirely without influence from the withheld epochs. The last step of the brain decoding mechanism is implemented using a nearest neighbour classifier between the predicted image feature vector *p_j_* from the EEG and the target image feature vector *t_j_*. This allows us to order all the images in our database (including the target stimuli) in the image feature space by their distance from the predicted feature vector. We can then convert this ordered list of stimuli into a rank by counting how far down the list the target image is, where a perfect prediction results in rank one and where the expected rank assuming chance performance is nS/2, where nS is the number of stimuli images.

## Results

### Datasets

Two different datasets are used to evaluate our zero-shot prediction architecture, in order to reduce the risk of overfitting to a particular dataset and limiting the generality of our conclusions. To facilitate comparison with previous approaches, two datasets with similar tasks, “Trento” and “Stanford”, are used.

### Trento Data

The first collection of EEG data analysed in this study is the Trento set [10] which uses 60 grayscale photographs as stimuli. Since this dataset was initially designed for classification, images are grouped in 30 pictures of 30 different land mammals and 30 pictures of 30 different hand tools. However, as explained in the introduction, in our image retrieval setting this category-level information is discarded and the stimuli is treated as 60 individual images. In an EEG experimental session, these images were each presented six times to the participant, for a total of 360 trials (i.e. 360 epochs). There were three participants, two of which took part in two experimental sessions and one participant who took part in one session. Participants were instructed to silently name the image with whatever term occurs naturally whilst EEG data was collected with a 64-channel EEG headset sampling at 500Hz. More details of the paradigm and recording of the data can be found in Murphy et al. [10]. The epoched data for each session therefore consists of a matrix of shape nE × nC × nT, where nE = 360, nC = 64 and nT = 500. Through the preprocessing steps outlined in the ‘Preprocessing Approach (FASTER)’ Section (including removal of noisy epochs), the resulting cleaned set was a matrix of size 340 × 7680 on average per recording. The number of epochs is approximate as for each experimental session, a different number of low quality epochs are removed during preprocessing. In the original study, the aim was to train a linear binary classifier to distinguish between epochs associated with mammal or tool stimuli, which differs from our goal of matching epochs to particular images. As such, the Trento materials use a narrow selection of stimuli from just two semantic categories, and each object image will be visually and semantically similar to many other images in the set. This provides a strong test of our methods ability to predict the correct image from a set of possible and very close alternatives.

### Stanford Data

The second EEG dataset we used to test our approach is an open dataset compiled at Stanford University [20]. Participants were presented with a series of colour photographs, drawn from the categories *human body, human face, animal body, animal face, fruit/vegetable*, or *man-made* (inanimate object). There were 12 images in each category and each image was presented 12 times in random order for a total of 864 trials per recording. Again, categories are discarded and the experiment is treated as an image retrieval task with 72 individual images. There were 10 participants, all of whom completed two sessions which each comprised of three separate EEG recordings for a total of 60 recordings. The EEG was recorded using a 128-channel headset sampling at 1kHz. Each recording therefore contained 864 epochs, each with 128,000 features in its raw form. The resulting cleaned set after preprocessing measured approximately 792 epochs × 128 channels × 120 timepoints, giving a EEG feature matrix of size 792 × 15,360 per recording. Across the recordings in the Stanford dataset, preprocessing resulted in the interpolation of approximately five channels, the removal of four independent components, and the removal of 0-2 trials of each stimulus. Following removal of noisy trials during preprocessing, four of these recordings were left with no trials for one of their stimuli; these recordings were excluded from further analysis.

### Measure of Accuracy

Given the difficulty of the zero-shot prediction task, we used an accuracy metric more sensitive to small improvements in prediction power based on the Cumulative Match Curve (CMC). Once a set of predicted visuo-semantic image features is produced for the EEG associated to a particular image presentation, all the images were ranked by their Euclidean distance from the predicted feature vector. A CMC was then generated by counting how often the true target appears in the top X ranked images as shown in Figure 4. For example, the first value on the x-axis represents the percent of cases where the target image is the nearest to the predicted features in the feature space, the second value on the x-axis represents the percent of cases in which the target image was one of the two closest images to the predicted features, and so on. The Area Under Curve (AUC) is calculated as the normalised volume below the curve for use as the final metric.

### Parameter Optimisation

A short gridsearch was performed to empirically optimise the parameters. A random recording from each dataset was chosen and used to perform this gridsearch for each experiment below. We then used the highest performing parameter set to perform the decoding for the rest of the recordings with the same dataset and image feature set. We do expect that different recordings will perform best under different parameter settings, and as such accuracy could be maximised with a more rigorous approach to gridsearching. That said we have chosen to determine parameters from a single recording in order to better reflect training in a real-world BCI system. In Tables 1 and 2 the recording used for each dataset has been marked (GS) and removed from the mean column.

**Table 1.** Trento Decoding Accuracies (CMC AUC %).

| Feature Set | S1_R1 | S2_R1 | S3_R1 | S1_R2 | S2_R2 | S4_R1 | GS | Mean |
| --- | --- | --- | --- | --- | --- | --- | --- | --- |
| Visual | 57.2 | 65.12 | 58.72 | 55.75 | 64.58 | 67.18 | 62.03 | 62.56 |
| Semantic | 57.74 | 64.74 | 56.26 | 61.66 | 65.98 | 63.9 | 64.28 | 61.71 |
| Visuo-semantic | 57.63 | 63.32 | 58.46 | 62.8 | 62 | 63.72 | 66.06 | 61.32 |

**Table 2.** Stanford Decoding Accuracies (CMC AUC %).

| Feature Set | S1 | S2 | S3 | S4 | S5 | S6 | S7 | S8 | S9 | S10 | GS | Mean |
| --- | --- | --- | --- | --- | --- | --- | --- | --- | --- | --- | --- | --- |
| Visual | 59.27 | 56.91 | 57.28 | 57.51 | 62.57 | 60.96 | 56.59 | 54.5 | 54.11 | 61.08 | 61.97 | 58.08 |
| Semantic | 60.12 | 56.89 | 61.38 | 60.35 | 65.49 | 63.93 | 62.36 | 55.93 | 58.07 | 64.28 | 64.82 | 60.88 |
| Visuo-semantic | 61.86 | 58.45 | 60.8 | 61.85 | 67.9 | 66.08 | 63.07 | 56.59 | 57.84 | 65.77 | 66.01 | 62.02 |

Alpha values {1e-2, 1e-1, 1e0, 1e1, 5e1, 1e2} were tested for the ridge regressor. The number of EEG features retained during feature selection (‘EEG Feature Selection’ Section) was tested over the values {25, 50, 75, 100, 125, 150, 175, 200, 250, 500, 750, 1000, 1500, 2000, 2500, 3000}.

### Decoding Accuracy

In order to compare the effectiveness of our chosen image feature models and confirm our expectation that combining the models would provide more predictive power than using them in isolation, the AUC for both datasets were calculated when using all visuo-semantic features (features_*vs*_) and compared against using only visual feature set (features_*v*_) or the semantic feature set (features_*s*_) individually.

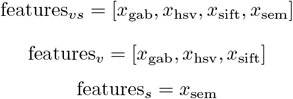

Results are shown in Table 1 for the Trento dataset and Table 2 for the Stanford dataset.

All the exemplar decoding results we present above are significantly above chance (50%), indicating a mapping between EEG activity and the image feature sets we have chosen that can be used for zero-shot brain decoding. For the Stanford dataset, which uses a larger and more diverse set of object images, the semantic feature set gives better accuracy than the visual feature set, and best performance is obtained with the combined visuo-semantic feature set. In the results for the Trento dataset, these trends are less clear, but it can be seen that the combination of all features is a robust approach overall.

### EEG Feature Selection Visualisation

In order to demonstrate that our EEG feature selection was performing as expected and was properly selecting activity from channels and timepoints known to relate to meaningful visual processing, we analysed which EEG features were assigned the highest score by the stability measure outlined in the ‘EEG Feature Selection’ Section. Figures 3a and 4a show a grand average of the stability scores at different times during an epoch. These values were generated by taking the mean of the scores across each channel for the time offset in question. Figures 3b-d and 4b-d show snapshots of the stability values at particular timepoints from the temporal plots, distributed over the EEG sensor locations.

**Fig 3.**
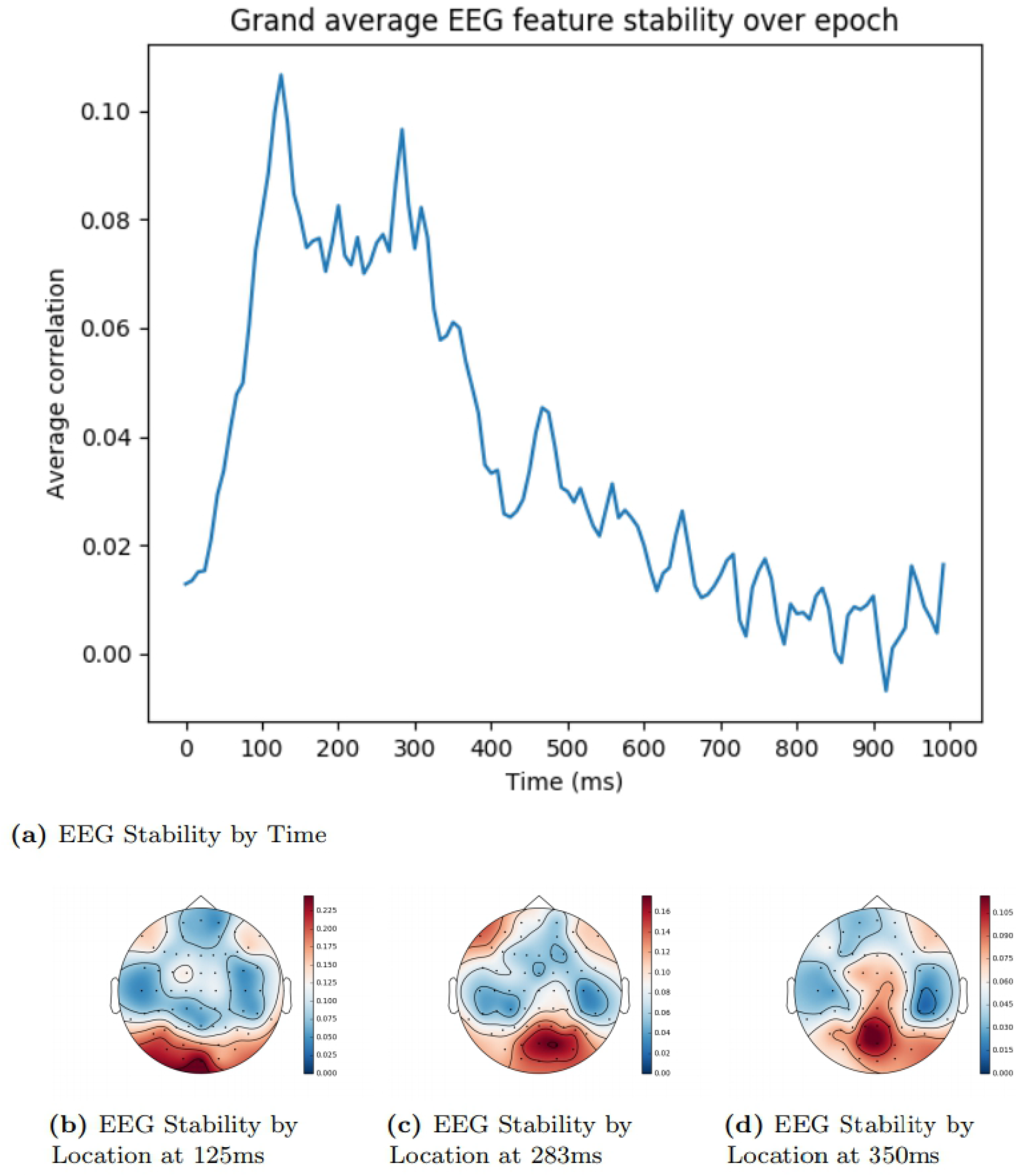
Trento EEG Feature Stability by Time and Location. The areas shaded in red signify the locations with highest EEG stability, while areas shaded in blue signify the lowest.

**Fig 4.**
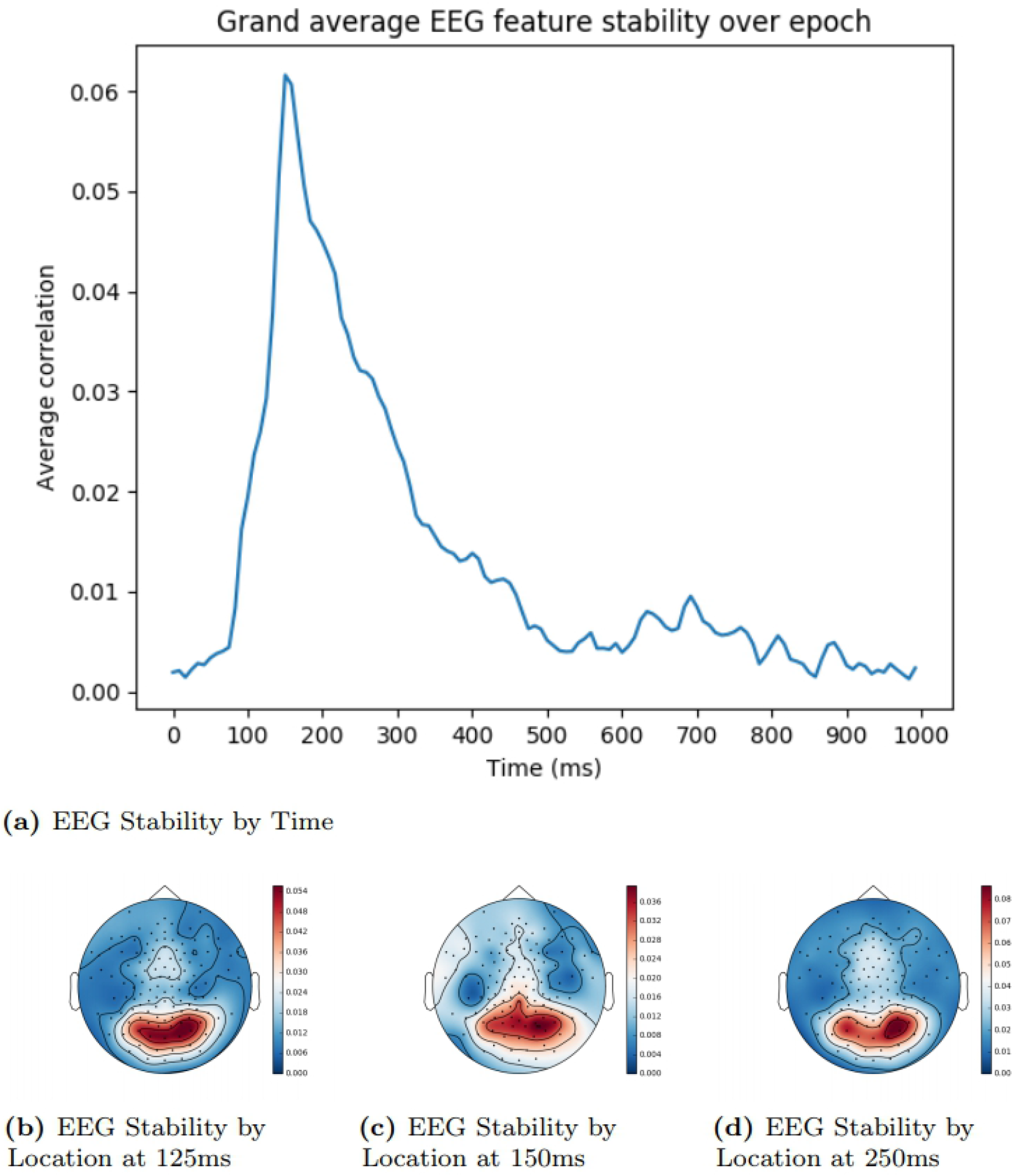
Stanford EEG Feature Stability by Time and Location. The areas shaded in red signify the locations with highest EEG stability, while areas shaded in blue signify the lowest.

Stability peaks within the expected time window [16], indicating that the feature selection method is properly determining the most informative features within the EEG activity. Based on previous research [16, 17, 27], we would expect the posterior sensors to be more useful early in an epoch when there is more visual processing, and that informative areas later on would be more spatially diffuse, when activity reflects more widely-distributed semantic processing of the stimulus. The stability analysis reflects this pattern, more clearly in the Trento dataset than the Stanford dataset.

### Comparison with State-of-the-art

Because of our zero-shot analysis framework, a study with directly comparable results could not be identified in a review of relevant EEG literature. However, the studies mentioned in the background section can provide a frame of reference. While Palatucci et al. [14] used image stimuli and decoded the image from brain activity, the focus was on decoding semantic information about the object in the image rather than retrieving the stimulus image based on the brain data. The datasets we have access to in this study involve much more visually complex image stimuli. Where Palatucci et al. [14] made use of minimilistic line drawings, the photographs used in both datasets analysed in this study are much more visually complex. In order to best leverage this extra visual information, we added several visual feature sets to our analysis.

The leave-one-class-out task performed by Palatucci et al. [14] is similar enough to the task in this study to give context to our results, though given the two studies use different datasets a direct comparison with our approach is not possible. The paradigm used in this study was very similar to those used in the Trento and Stanford experiments, with participants being presented with a series of images and asked to silently name them. Compared with the Palatucci et al. [14] study, we obtain slightly stronger results (Table 3).

**Table 3.** Leave-one-class-out task percent rank accuracy.

| Dataset Name | Rank Accuracy |
| --- | --- |
| Palatucci et al. [14] (their approach) | 56% |
| Trento dataset (our approach) | 61.1% |
| Stanford dataset (our approach) | 61.65% |

## Conclusion

In this paper we proposed an approach to zero-shot image retrieval in EEG data using a novel combination of feature sets, feature selection, and regression modeling. We have shown that a combination of visual and semantic feature sets performs better than using either of those feature sets in isolation. We also analysed the performance of each image feature model used in our approach individually to help identify where future improvements could be made.

We hope that future work can improve upon this approach using the same open dataset for comparison as it is difficult to accurately predict how well our approach would perform on the other datasets mentioned in Sections ‘Introduction’ and ‘Comparison with State-of-the-art’. We have demonstrated that the features we extracted for the EEG data and images are justified and perform significantly above chance. However it is possible that our image features do not accuracy reflect all stages of human visual processing, and that a different set of features would better facilitate a regression model. For example, large neural networks that recognise images have a hierarchical architecture which reflects some aspects of human visual processing [42, 43] and which could provide an effective model for our feature space. Alternatively we could replace our semantic features with a set derived from a distributional word embedding model such as word2vec [44] or fastText [45].

Moreover, our EEG feature selection may correctly quantify the usefulness of each particular timepoint in each channel, however it is likely that features which are close in time and location will have very similar information and thus similar scores, and so a feature selection method may select a set of good quality but redundant features. In future work, we will explore feature selection methods that produce a small set of maximally informative EEG features. Nevertheless, our approach has demonstrated a marked improvement over current state-of-the-art for EEG zero-shot image decoding and is a step towards the application of EEG to real-world BCI technologies.

